# ViralConsensus: A fast and memory-efficient tool for calling viral consensus genome sequences directly from read alignment data

**DOI:** 10.1101/2023.01.05.522928

**Authors:** Niema Moshiri

## Abstract

**Motivation:** In viral molecular epidemiology, reconstruction of consensus genomes from sequence data is critical for tracking mutations and variants of concern. However, as the number of samples that are sequenced grows rapidly, compute resources needed to reconstruct consensus genomes can become prohibitively large.

**Results:** ViralConsensus is a fast and memory-efficient tool for calling viral consensus genome sequences directly from read alignment data. ViralConsensus is orders of magnitude faster and more memory-efficient than existing methods. Further, unlike existing methods, ViralConsensus can pipe data directly from a read mapper via standard input and performs viral consensus calling on-the-fly, making it an ideal tool for viral sequencing pipelines.

**Availability:** ViralConsensus is freely available at https://github.com/niemasd/ViralConsensus as an open-source software project.

**Supplementary information:** Supplementary data are available online.

## 1 Introduction

Viral molecular surveillance, a technique in which viral genomes are reconstructed from sequence data generated by samples collected from patients as well as the environment (e.g. wastewater) and are monitored in real-time or near real-time, has been critical throughout the COVID-19 pandemic (Oude Munnink *et al*., 2021; Karthikeyan *et al*., 2022). The reconstruction of consensus genome sequences from raw sequence data requires the use of various bioinformatics pipelines, which can be slow and can require non-trivial computational expertise (Truong Nguyen *et al*., 2021; Posada-Céspedes *et al*., 2021; Moshiri *et al*., 2022).

The current best-practice pipeline for reconstructing a consensus genome sequence from raw viral amplicon sequence data is the iVar pipeline (Grubaugh *et al*., 2019). First, reads are mapped to the reference genome using a read mapper such as Minimap2 (Li, 2018) or BWA (Li & Durbin, 2009) and position-sorted using samtools (Li *et al*., 2009). Next, reads are primer- and quality-trimmed using iVar and again position-sorted using samtools. A pile-up is then computed from the sorted trimmed reads using samtools, and a consensus genome sequence is called from the pile-up file using iVar. This position-sorted pile-up-based approach is ideal for long genomes (e.g. human) in which the memory needed to store counters for every position of the genome simultaneously would become prohibitively large, but due to their small length, viral consensus genome sequences can be computed much faster.

Here, we introduce ViralConsensus, a fast and memory-efficient tool for calling viral consensus genome sequences directly from read alignment data. ViralConsensus is orders of magnitude faster and more memory-efficient than existing methods. Further, unlike existing methods, ViralConsensus can pipe data directly from a read mapper via standard input and performs viral consensus calling on-the-fly, making it an ideal tool for viral sequencing pipelines.

## 2 Results and discussion

ViralConsensus is a command-line tool written in C++ and depends on htslib (Bonfield *et al*., 2021). ViralConsensus takes the following as required input: (i) a SAM/BAM/CRAM file containing the mapped reads (or “-” to read from standard input), (ii) a FASTA file containing the reference genome, and (iii) an output FASTA file to write the consensus genome (or “-” to write to standard output). Optionally, the user can also provide the following: (i) an output file in which to write base counts at each position (or “-” to write to standard output), (ii) an output file in which to write the insertion counts (or “-” to write to standard output), (iii) a minimum quality threshold to count a base in a read (default: 20), (iv) a minimum depth to call a position in the consensus (default: 10), (v) a minimum frequency to call a base/insertion in the consensus (default: 0.5), (vi) a symbol to use for ambiguous positions (default: “N”), (vii) a BED file containing primer positions to trim (default: no primer trimming), and (viii) a number of positions beyond the end of a primer to also trim (default: 0).

ViralConsensus first loads the reference genome and preallocates base/insertion counters for each position of the genome, and it then streams the reads on-the-fly without any need for sorting, and it increments the base/insertion counts at each position covered by the reads. ViralConsensus is able to trim reads on-the-fly using the user-provided minimum base quality threshold (for quality-trimming) and primer file (for primer-trimming), but users can trim reads prior to executing ViralConsensus if desired. Because it performs all computations on-the-fly and does not require intermediary files, ViralConsensus can be easily integrated into existing pipelines by piping directly from the read mapper, significantly reducing disk I/O. Further, because of its approach, ViralConsensus has constant memory consumption and linear runtime with respect to sequencing depth, and it has linear memory consumption with respect to genome length.

In order to benchmark ViralConsensus with respect to sequencing depth, we obtained a SARS-CoV-2 amplicon sequencing dataset in which 2,607 samples were sequenced PE150 across four lanes of an S4 flow cell to an average read count of 4.58 M read pairs per sample using the SWIFT v2 protocol on an Illumina NovaSeq 6000 (Moshiri *et al*., 2022). Samples were mapped to the NC_045512.2 reference genome using Minimap2. We selected the single highest-depth sample and randomly subsampled it to *n* = 100, 1K, 10K, 100K, and 1M successfully-mapped reads, with 10 replicates for each *n*. We then ran ViralConsensus as well as the iVar pipeline to compute consensus sequences from each subsampled replicate.

As can be seen in Figure 1, ViralConsensus is orders of magnitude faster and more memory efficient than the iVar pipeline, and it is able to call a consensus sequence from an amplicon sequencing dataset with 1 million reads in less than 2 seconds with a peak memory usage of less than 12 MB. Importantly, while both methods’ runtimes scale linearly with sequencing depth, the iVar pipeline’s peak memory usage grows substantially as sequencing depth increases, whereas the peak memory of ViralConsensus remains constant.

**Fig. 1.**
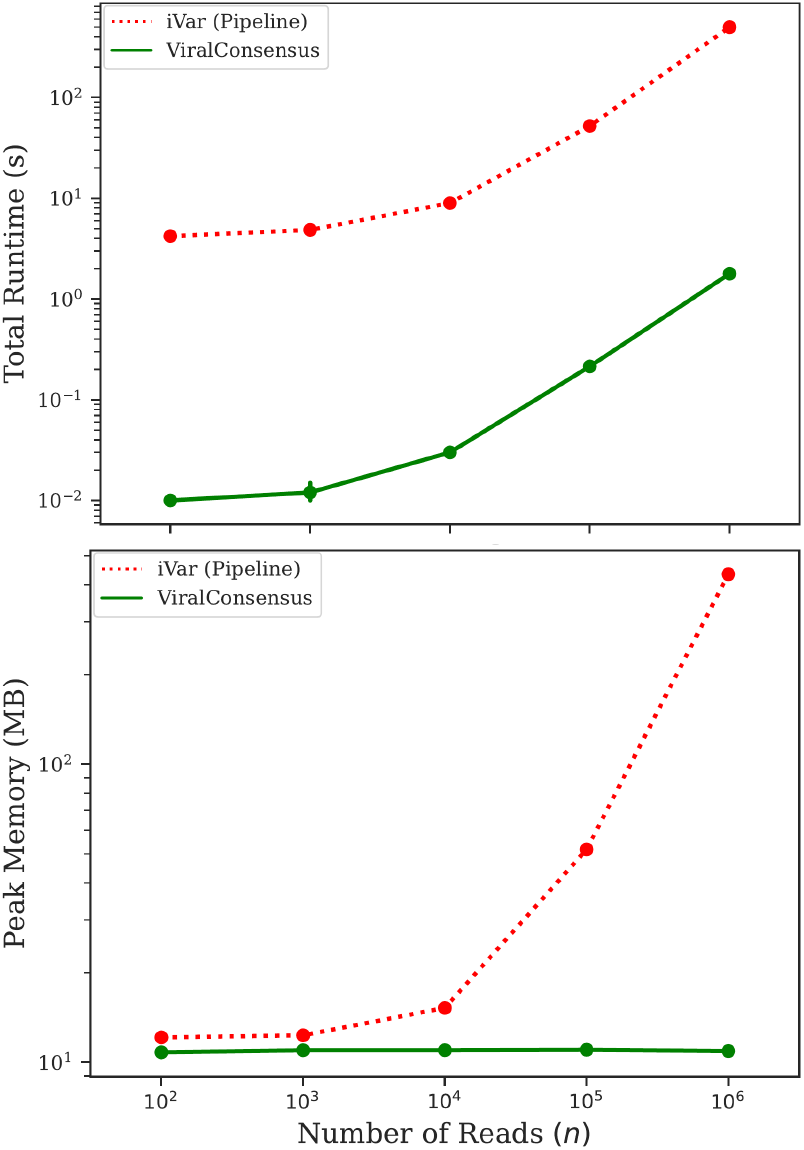
Benchmark. Total runtime (top) and peak memory (bottom) for SARS-CoV-2 sequence datasets with *n* = 100, 1K, 10K, 100K, and 1M mapped reads. All runs were executed sequentially on a 2.8 GHz Intel i7-1165G7 CPU with 16 GB of memory.

## Acknowledgements

We would like to thank Tajana Rosing, Behnam Khaleghi, Josh Cross, Ronak Shah, Niya Shao, Sean Eilert, Ameen Akel, Justin Eno, Ken Curewitz, Rob Knight, Kristian Andersen, and Karthik Gangavarapu for fruitful conversations.

## References

Bonfield, J.K. et al. (2021) HTSlib: C library for reading/writing high-throughput sequencing data. GigaScience, 10(2), giab007.

Grubaugh et al. (2019) An amplicon-based sequencing framework for accurately measuring intrahost virus diversity using PrimalSeq and iVar. Genome Biol., 20(1), 8.

Karthikeyan, S. et al. (2022) Wastewater sequencing reveals early cryptic SARS-CoV-2 variant transmission. Nature, 609, 101–108.

Li, H. (2018) Minimap2: pairwise alignment for nucleotide sequences. Bioinformatics, 34(18), 3094–3100.

Li, H. & Durbin, R. (2009) Fast and accurate short read alignment with Burrows–Wheeler transform. Bioinformatics, 25(14), 1754–1760.

Li, H. et al. (2009) The Sequence Alignment/Map format and SAMtools. Bioinformatics, 25(16), 2078–2079.

Moshiri, N. et al. (2022) The ViReflow pipeline enables user friendly large scale viral consensus genome reconstruction. Sci. Rep., 12, 5077.

Oude Munnink, B.B. et al. (2021) The next phase of SARS-CoV-2 surveillance: real-time molecular epidemiology. Nat. Med., 27, 1518–1524.

Posada-Céspedes, S. et al. (2021) V-pipe: a computational pipeline for assessing viral genetic diversity from high-throughput data. Bioinformatics, 37(12), 1673–1680.

Truong Nguyen, P.T. et al. (2021) HAVoC, a bioinformatic pipeline for reference-based consensus assembly and lineage assignment for SARS-CoV-2 sequences. BMC Bioinf., 22, 373.

